# Continuum theory for the mechanics of curved epithelial shells by coarse-graining an ensemble of active gel cellular surfaces

**DOI:** 10.1101/2025.02.27.640501

**Authors:** Pradeep K. Bal, Adam Ouzeri, Marino Arroyo

## Abstract

Epithelial tissues undergo complex morphogenetic transformations driven by cellular and cytoskeletal dynamics. To understand the emergent tissue mechanics resulting from sub-cellular mechanisms, we formulate a fully nonlinear continuum theory for epithelial shells that coarse-grains an underlying 3D vertex model, whose surfaces are in turn patches of active viscoelastic gel undergoing turnover. Our theory relies on two ingredients. First, we relate the deformation of apical, basal and lateral surfaces of cells to the continuum deformation of the tissue mid-surface and a thickness director field. We explore two variants of the theory, a Cosserat theory accommodating through-thickness tilt of cells, and a Kirchhoff theory assuming that lateral cell surfaces remain perpendicular to the mid-surface. Second, by adopting a variational formalism of irreversible thermodynamics, we construct an effective Rayleighian functional of the tissue constrained by the cellular-continuum kinematic relations, which therefore depends on continuum fields only. This functional allows us to obtain the governing equations of the continuum theory and is the basis for efficient finite element simulations. Verification against explicit 3D cellular model simulations demonstrates the accuracy of the proposed theory in capturing epithelial buckling dynamics. Furthermore, we show that the Cosserat theory is required to model tissues exhibiting apicobasal asymmetry of active tension. Our work provides a general framework for further studies integrating refined subcellular models into continuum descriptions of epithelial mechanobiology.

## 1. Introduction

Epithelial tissues control key morphogenetic events. Reshaping of epithelial sheets relies on cellular-level mechanisms such as cell division, apoptosis or cell rearrangements [1, 2], on actively generated forces at sub-cellular scales, leading for instance to active bending by apicobasal asymmetries of contractility [3, 4, 5, 6], on buckling following lateral compression [7, 8], or on a combination of the above [9, 10, 11]. In recent years, the study of the impact of cellular and cytoskeletal dynamics on epithelial tissue reshaping has been enabled by engineered in vitro systems offering a high degree of control and accessibility. Such systems include organoids [9, 6, 12], suspended [13, 14, 15] or encapsulated [16] cell monolayers, optogenetically stimulated epithelial sheets [17, 18], or pressurized epithelial domes [19]. These experimental advances, enabling the characterization of cell and tissue shape while monitoring and manipulating the dynamics of the cytoskeleton, call for mechanistic models of the active mechanics of epithelial tissues that connect sub-cellular and mesoscopic scales.

Three-dimensional vertex models provide a direct connection between possibly heterogeneous cellular properties, cellular shape, cell rearrangements and tissue shape [20, 21, 9, 5, 2, 12]. Epithelial reshaping has also been studied with continuum surface models, which provide a coarse-grained description of the cellular tissue, are amenable to mathematical analysis, and may enable more efficient simulations [22, 3, 23, 24, 25, 26]. Interestingly, these models raise questions about the role of discreteness and cellularity in the context of different morphogenetic processes. Continuum models are generally based on balance laws supplemented with phenomenological constitutive relations. For instance, [3] adapted a conventional Koiter model for thin elastic shells and introduced apicobasal asymmetry through a spontaneous curvature parameter, whereas [8] used a 3D continuum approach to model an asymmetric and growing epithelial layer. Instead, [24, 25] developed and applied a general phenomenological theory for active surfaces. One issue with this traditional approach, particularly when applied to epithelial shells, is the large number of active, dissipative and elastic model parameters, which cannot be unambiguously estimated with current experiments or related to cellular mechanism. One alternative is to derive continuum models by coarse-graining or homogenizing smaller-scale descriptions of the sub-units controlling the behavior of the supra-cellular assembly. Arguably, for epithelial monolayers these sub-units are the actomyosin gels forming the cortical surfaces, which enclose roughly constant cellular volumes [27], other cytoskeletal structures such as intermediate filament networks [19, 13, 28, 29], and cell-cell and cell-matrix adhesions. A direct approach to coarse-graining, widely used to homogenize atomistic models into continuum mechanics theories [30, 31, 32] and also applied to epithelial sheets [33], relies on relating the deformation of the continuum surface and that of the sub-units through the so-called Cauchy-Born rule; such an kinematic approach does not introduce additional material parameters other than those of the microscale model. In the context of epithelial mechanics, this approach could enable the systematic integration of refined and experimentally informed models of cytoskeletal dynamics into continuum models of the tissue.

Here, we formulate a continuum surface model for epithelial shells based on an underlying 3D cellular (vertex) model, whose surfaces are in turn patches of active viscoelastic gel undergoing turnover [34]. See Figure 1 for an illustration of the proposed approach. The resulting model is described by a parametrized surface, describing the overall shape of the cell sheet, along with a thickness director field, dynamically adjusted in agreement with the assumption of constant cellular volume and allowing us to define apical and basal surfaces as offsets. This theory provides relations between the continuum fields and the deformation of apical, basal, and lateral surfaces within the cell monolayer. This kinematic connection between the continuum and the cellular deformations allows us to integrate the collective effect of all cortical cellular faces into a homogenized continuum model, which inherits the active and viscoelastic character of the microscopic active gel model [35]. A variational formulation of irreversible thermodynamics facilitates a direct homogenization procedure from cortical active gels to an effective tissue model.

**Figure 1:**
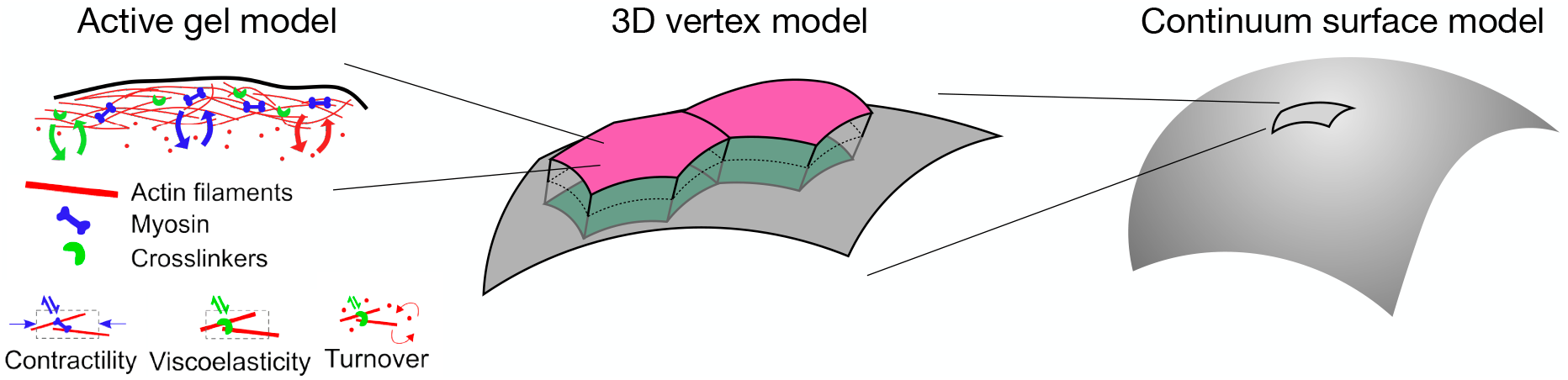
Illustration of the proposed approach to model the mechanics of epithelial shells.

A few previous works have developed homogenized continuum models from 2D vertex models of epithelial tissues. Focusing on fluid-like tissues, a connection has been made between continuum hydrodynamics and cellular events including topological transitions of the junctional network [36, 37]. Here, we focus in solid-like tissues without topological transitions, either because they are in a jammed state or because the observation times are relatively short. In this limit, [33] proposed an effective surface model obtained by coarse-graining a 2D vertex model of apical surfaces. However, the bending resistance of the resulting model does not arise from through-the-thickness moments. In [38, 23], a 2D lateral vertex model was coarse-grained to obtain effective elastica models, capturing effects depending on apicobasal asymmetry [2]. This kind of approach, constrained to spherical cap geometries, was applied to study the morphology of intestinal organoids [6]. Akin to Kirchhoff-Love theories of thin shells, these models assume that lateral edges remain normal to the mid-line curve. This assumption is usually valid when shells are very thin, but epithelial monolayer, particularly columnar ones, can exhibit deformation features comparable to their thickness. More recently, a continuum theory coarse-graining lateral vertex models and accounting for through-the-thickness shear, akin to a Cosserat theory of shells, has been developed [26]. None of these models account for time-dependent behaviors stemming from cytoskeletal dynamics, nor coarse-grain a 3D vertex model into a continuum surface.

The paper is organized as follows. In Section 2., we develop a mathematical model for an active and viscoelastic cortical surface undergoing turnover. This includes a review of the kinematics of a generic continuous material surface, pertinent to both individual cellular faces and to the tissue, Section 2.1, and a summary of the active gel model used here, Section 2.2. In Sections 3. and 4., we introduce the Cosserat and Kirchhoff theories for epithelial shells. Section 5. particularizes the modeling framework to specific constitutive choices, and identifies the fundamental set of parameters of the cell monolayer that control its mechanics. The finite element implementation of the theory is briefly discussed in Section 6.. We exemplify the application of the theory in Section 7., where we perform a comparison between the proposed model for epithelial shells and a detailed 3D cellular model. Finally, we collect our conclusions and outline the limitations and possible extensions of the proposed model in Section 8..

## 2. Modeling a cortical surface

### 2.1 Surface kinematics

We summarize the mathematical description of the kinematics of deforming surfaces, without any assumption about the smallness of deformations. We refer the reader to [39, 40, 31, 41] for further background and to Figure 2 for an illustration. Consider a time-evolving continuous material surface Γ_*t*_, which here represents a cortical surface within a cell, but later can represent the mid-surface of an epithelial sheet. We parametrize this surface with a time-dependent map from a planar parametric domain 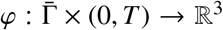 such that 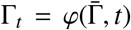. We describe both 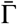 and ℝ^3^ Cartesian coordinates, { *ξ*^1^, *ξ*^2^} and {*x*^1^, *x*^2^, *x*^3^} . We introduce the natural tangent vectors to the surface induced by the parametrization ***g***_*I*_ = *∂φ*/*∂*ξ^*I*^, and the unit normal to the surface

**Figure 2:**
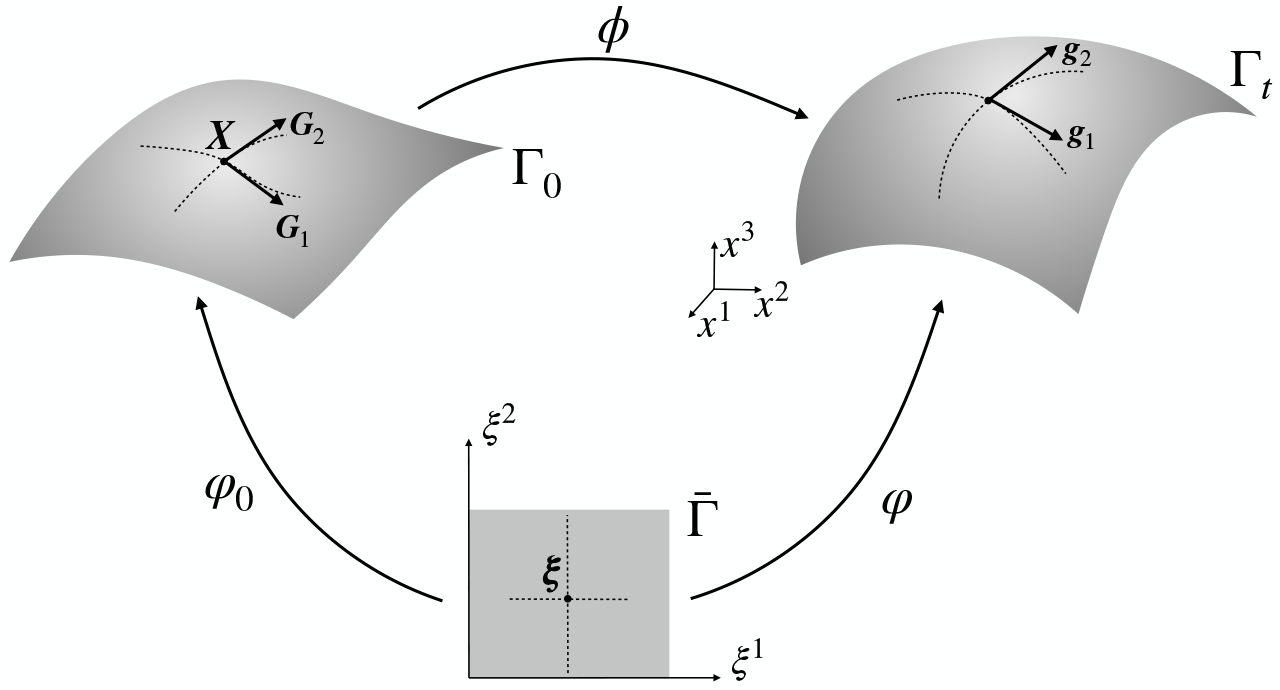
Illustration of the kinematics of a deforming material surface.

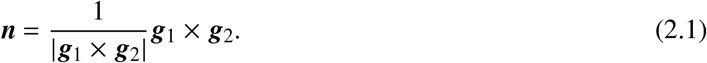

The metric tensor of Γ_*t*_ induced by the Euclidean metric of ℝ^3^ is *g*_*IJ*_ = ***g***_*I* ·_ ***g***_*J*_. Its inverse is defined by the relations 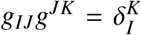. Here and elsewhere, repeated indices imply summation, from 1 to 2 for uppercase indices and from 1 to 3 for lowercase indices. Introducing 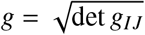, the area element of the surface can be expressed as *dS* = *g dξ*^1^*dξ*^2^. The coefficients of the second fundamental form, describing the curvature of the surface, can be computed as

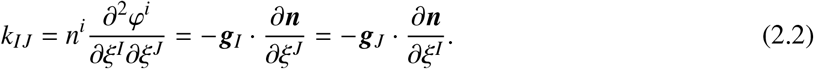

We consider an arbitrary reference configuration of the material surface, described by the map 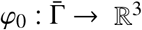 such that 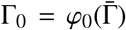. We can also define the natural tangent vectors ***G***_*I*_ = *∂φ*_0_/*∂*ξ^*I*^ and metric tensor of this reference state as *G*_*IJ*_ = ***G***_*I*_ · ***G***_*J*_. Analogously, we introduce its inverse defined by the relations 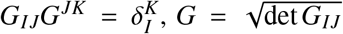, the area element of the surface *dS* _0_ = *G d*ξ^1^*d*ξ^2^, and denote the second fundamental form of Γ_0_ by ***K***. We assume that the map *φ*_0_ is a fixed diffeomorphism, and therefore we can identify functions defined over the parametric or the reference domains.

The deformation map is defined as 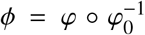. The deformation gradient is the derivative of this map ***F*** = *Dϕ* = *Dφ* ° (*Dφ*_0_)^−1^, where here ^−1^ should be interpreted as the pseudoinverse of a 3 × 2 matrix. The Lagrangian velocity field is given by 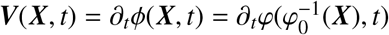 with ***X*** ∈ Γ_0_, and the Eulerian velocity by ***v*** = ***V*** °*ϕ*^−1^. The right Cauchy-Green deformation tensor is a symmetric positive definite tensor field defined in the tangent bundle of Γ_0_ characterizing the local in-plane deformation of the surface. In the natural basis {***G***_1_, ***G***_2_}, its components are simply *C*_*IJ*_ = *g*_*IJ*_. This tensor allows us to compute the stretch ratio along a direction in the reference configuration Γ_0_, specified by a unit tangent vector ***M*** = *M*^*I*^***G***_*I*_ with *G*_*IJ*_ *M*^*I*^ *M*^*J*^ = 1 , as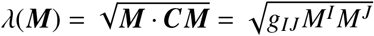 . The components of vector ***M*** in the basis {***G***_1_, ***G***_2_} can be computed as *M*^*I*^ = *G*^*IJ*^(***M* · *G***_*J*_). The invariants of ***C*** are naturally expressed in terms of its mixed components *C*^*I*^ _*J*_ = *G*^*IK*^*g*_*KJ*_. For instance, the Jacobian determinant measuring the local area change is given by 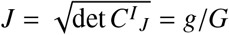.

### 2.2 Active viscoelastic gel for cellular faces

We summarize next the active viscoelastic gel model for the actin cortex considered here. We formulate this model for a generic cortical patch, which can represent an apical, a basal or a lateral face of the tissue. At short times, this material behaves as a prestressed crosslinked network of semi-flexible filaments, which we model with a hyperelastic membrane model with active tension. At longer time-scales, remodeling of the network by crosslinker and filament turnover should dissipate the elastic stresses. We account for viscoelastic relaxation by considering a time-dependent material metric tensor, which can be interpreted as a resting strain. The resulting model is similar to previous viscoelastic active gel fluids [35], with the difference that ours has nonlinear hyperelasticity as its short-time limit, and therefore the work done by closed deformation paths is zero by construction. This is not the case in general for models based on an underlying hypoelastic model, in which elasticity may result in uncontrolled power input or dissipation [42]. Our modeling framework can be adapted to the microscopic cytoskeletal model of choice.

To aid in the homogenization procedure, we adopt a variational formalism of irreversible thermodynamics, sometimes called Onsager’s variational principle, applicable to isothermal and low Reynolds systems. According to this formalism, emerging as a limit of bracket formalisms [43, 44], the dynamics of the system minimize a Rayleighian functional where energy-release-rate, dissipation and power input compete [45, 46, 47, 48, 49, 50, 51, 41].

#### Mass balance

We model the active gel as a continuous surface with a field of areal cortical density ρ, which if the three-dimensional density of the cortex is uniform can be interpreted as its thickness. Density is controlled by both deformation and turnover. We describe mass balance of the cortical layer as [35]

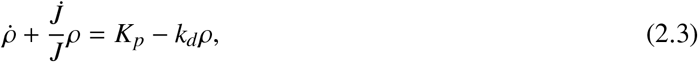

where the dot denotes material time differentiation, that is time differentiation at fixed ***X***. The second term in the left-hand-side models changes of density due to deformation, whereas the right-hand-side models turnover, with *K*_*p*_ the polymerization rate and *k*_*d*_ the depolymerization rate. According to this equation, the steady-state density is 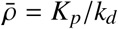. The cortical density per unit reference area can be computed as *ϱ*_0_ = *J*ρ. Mass balance can be expressed in terms of the reference density as

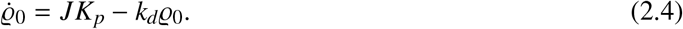

#### Stored elastic energy

To model the short-term elasticity of the cortical network, which may involve very large deformations, we consider a hyperelastic energy density per unit mass Ψ. According to the principle of objectivity, this function can be expressed in terms of ***C*** and ***G***, which we simply denote by Ψ(***C***) because once the reference configuration is fixed, so is ***G*** [40]. If the material is isotropic, then Ψ depends on deformation through the scalar invariants of right Cauchy-Green deformation tensor. In 2D, two invariants are sufficient, and we choose *I*_1_ = trace ***C*** = *C*_*IJ*_*G*^*IJ*^ = *g*_*IJ*_*G*^*IJ*^ and *I*_3_ = det ***C*** = det *g*_*IJ*_/ det *G*_*IJ*_ = *J*^2^. A general approach to model isotropic viscoelasticity in a finite deformation regime is to view the material metric as a dynamical variable [52, 53, 54, 55, 56], which we denote by G (***X***, *t*). Then, the hyperelastic potential is additively split into one component that is not susceptible to viscoelastic relaxation, denoted by 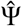, and another one susceptible to viscoelastic relaxation, denoted by 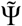, which depends on the invariants of ***C*** relative to this evolving metric, that is 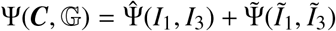 with

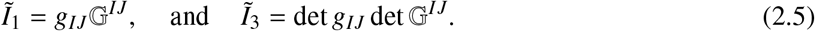

The material metric G is symmetric and positive definite, and can be interpreted as a tensorial quantity generalizing the idea of a time-dependent resting length [13]. The assumption that the actomyosin cortex can fully relax its elastic energy and become a fluid at long times is expressed by 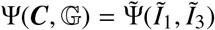 while the potential 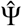 can be interpreted as modeling residual elasticity remaining in the cortical surfaces at long times. Physically, it can be ascribed to load-bearing structures other than the actomyosin cytoskeleton. For instance, the plasma membrane can become taut under large stretches, the adhesion molecules can become crowded and resist excessive deformation, and intermediate filaments accumulate at cellular surfaces and turnover slowly. All of these elements may prevent complete relaxation of elastic stresses at the cell surface, and hence provide residual elasticity.

Thus, the elastic energy of a cortical patch with reference domain 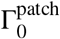 and deformation map *ϕ*^patch^ is a functional of the deformation, the evolving material metric and density

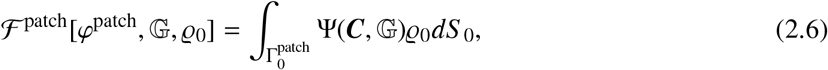

where we have written ℱ^patch^ as a functional of *φ*^patch^ rather than of 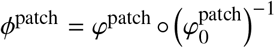 because 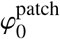 is fixed. Its rate of change can be expressed as

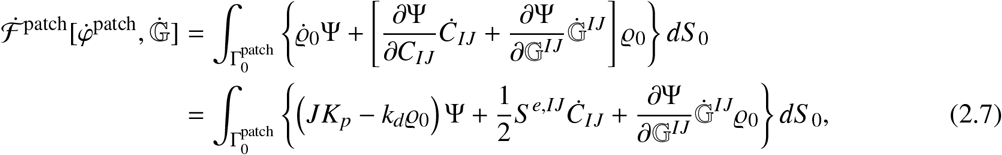

where we have introduced the equation of mass balance as a constraint and introduced the elastic second Piola-Kirchhoff stress tensor ***S***^*e*^ = 2ϱ_0_*∂*Ψ/*∂****C***. The elastic Cauchy or true stress tensor is therefore *σ*^*e*^ = (1/*J*)***FS***^*e*^ ***F***^*T*^ .

#### Dissipation

We also introduce a dissipation potential per unit mass that models the rate-dependent frictional resistance to changes in the material metric, 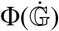 which in general can depend on deformation and material metric.

Consistency with the entropy production inequality demands that this function be convex in 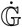 and attain its minimum at 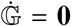 For concreteness, we can consider a quadratic dissipation potential independent of deformation and material metric of the form [54, 57]

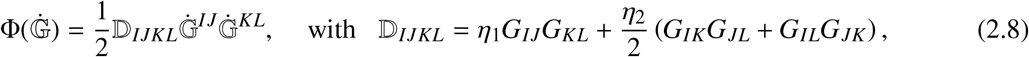

where D is a general isotropic viscosity tensor and *η*_1_ and *η*_2_ are viscosity parameters.

In addition to the viscous dissipation embodied in Φ, we consider the dissipation caused by the viscous drag associated to the surrounding medium, which we simplify as a local quadratic drag with coefficient 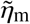, leading to the dissipation functional

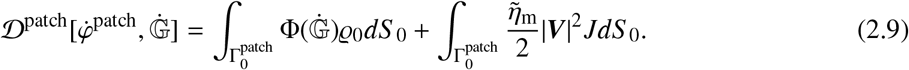

#### Active tension

We finally introduce the power input density modeling active contractility. One possibility is to introduce an active stress conjugate to G, which acts as an entropy sink and drives the evolution of the material metric away from the passive viscous relaxation. Instead, due to turnover of myosin motors, we consider that active tension acts isotropically in the Eulerian configuration of the material and consider a conventional power input per unit mass of the form ξ trace 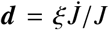 with ξ an activity parameter [58, 35], leading to a power input functional of the form

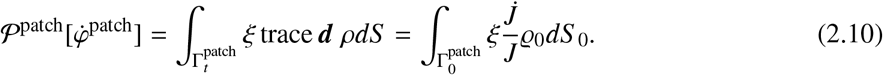

#### Governing equations

Onsager’s variational formalism of irreversible thermodynamics hinges on the formulation of the Rayleighian functional, which is minimized with respect to the generalized velocities to derive the governing equations. The variational structure enables a straightforward treatment of the kinematic constraints typical of homo-geneization procedures, as developed later. We introduce the system Rayleighian functional as

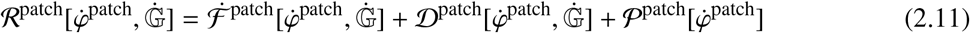

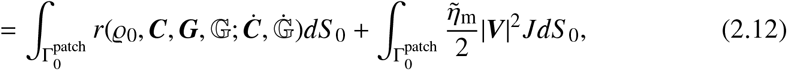

where we have introduced the Rayleighian density per unit reference area of the material surface

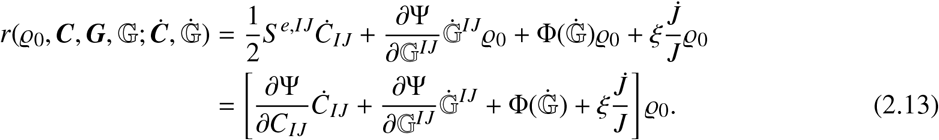

We note that we have ignored the first term in Eq. (2.7) because it does not contribute to the variational principle. The governing equations result from minimizing this functional with respect to 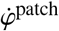 and to 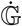 . Because ℛ^patch^ does not involve gradients in 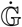, the corresponding stationarity conditions are systems of ordinary differential equations involving derivatives with respect to time, but not space, at each point in 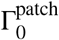. These equations specify the time evolution of the material metric and are given by [52]

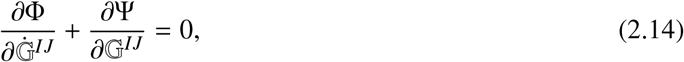

or more specifically for a quadratic dissipation potential by

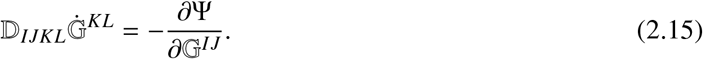

We also need to specify initial conditions for G. To model a situation in which the system is initially in an elastically relaxed state, the appropriate initial conditions are

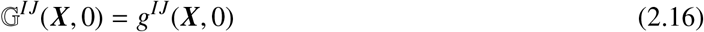

for all ***X*** in Γ_0_, since in this case 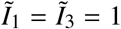

For interpretation purposes, we note that when 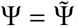, then it is a function of *C*_*IK*_ G^*KJ*^ only, and therefore by the chain rule it is possible to express *∂*Ψ/*∂* G^*KJ*^ in terms of *∂*Ψ/*∂C*_*IK*_. Using this fact, we can rewrite this kinetic law as

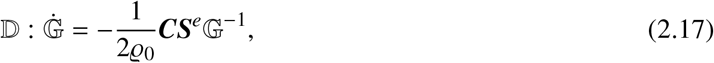

which shows that the elastic stress tensor drives the viscous evolution of the material metric.

The stationarity condition with respect to 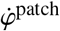 leads to the weak form of balance of linear momentum, normal and tangential in the surface, expressed as

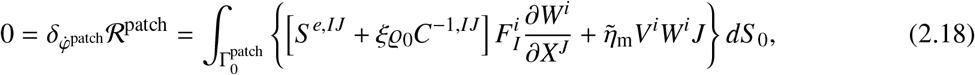

for all admissible variations ***W***. This expression allows us to identify the second Piola-Kirchhoff stress tensor for the surface as

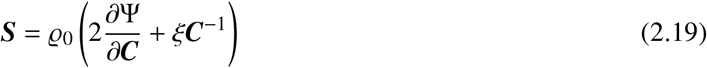

This stress tensor is 2D, and has therefore units of surface tension. The corresponding Cauchy stress tensor is *γ* = *γ*^*e*^ + *ξρ* ***g***. As further elaborated in [34], the strong form of the balance of linear momentum expressed in the Eulerian configuration takes the form

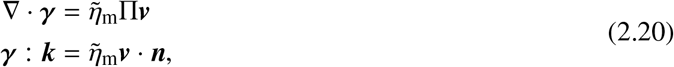

where ∇and Π are the covariant derivative and the tangential projection operator on Γ_*t*_. The first of these equations expresses tangential balance of linear momentum, and the second balance of linear momentum normal to the surface. The coupled governing equations for the density field, the material metric, and the velocity field (allowing us to evolve Γ_*t*_) are given by Eqs. (2.3,2.14,2.20).

## 3. Cosserat continuum theory for epithelial shells accounting for tilt

We start by introducing a general theory for epithelial shells allowing for through-the-thickness shear, which in the present context implies tilt of lateral surfaces. This theory can be viewed as a Cosserat theory [59], where the state of the shell is given not only by the parametrization of the mid-surface, but also by a director field describing how through-the-thickness material elements deform. In a related context, continuum theories of lipid bilayers extending the classical Helfrich model to account for lipid tilt have been developed to better describe lipid configurations close to membrane insertions [60, 61].

### 3.1 Cosserat kinematics of the tissue

The state of the system is described by the map of the mid-surface *φ*(***ξ***, *t*) and by vector field 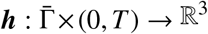, such that ±***h***/2 is the displacement of a point in the apical/basal surface relative to a corresponding point in the mid-surface. This additional vector field is not necessarily aligned with the normal to Γ_*t*_ to encode the tilt of lateral surfaces. Unlike common Cosserat shell theories, it is not a unit vector to further encode the thickness of the tissue. See Fig. 3 for a graphical illustration. Depending on the system and its coupling with a substrate, it may be convenient to parametrize the basal surface with *φ* and view ***h*** as the displacement of a point in the apical surface relative to the basal surface.

**Figure 3:**
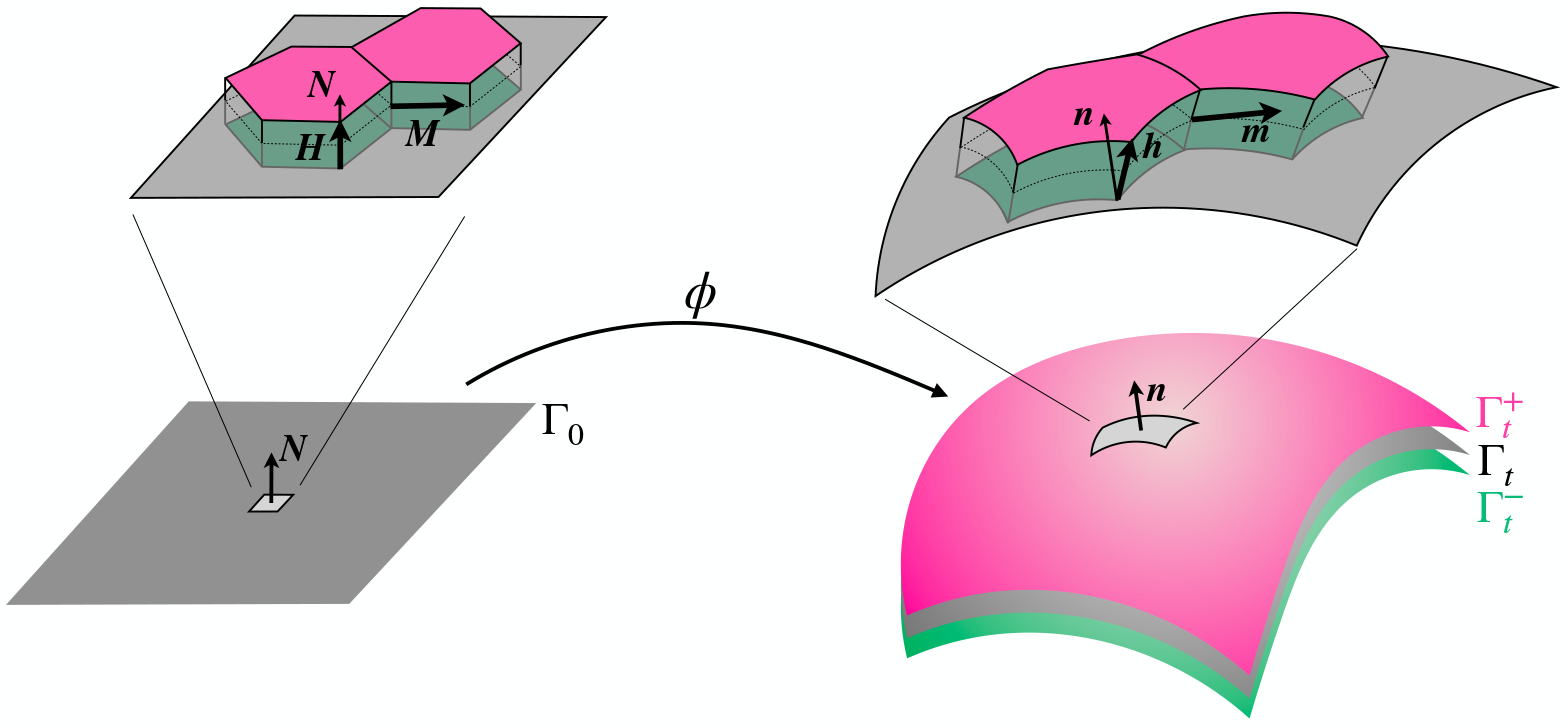
Illustration of the active bilayer Cosserat model accounting for tilt. For simplicity of illustration, we consider a planar reference configuration with normal reference field ***H*** (left). The through-the-thickness deformation and the thickness are encoded by the vector field ***h***, which is not necessarily aligned with the normal ***n***.

With the parametrization of the mid-surface and the director field, we can define the extended deformation map from a reference slab to the tissue layer as

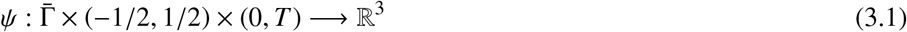

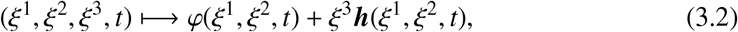

and an analogous map for the fixed reference configuration

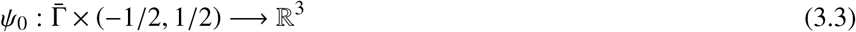

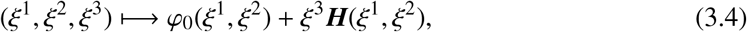

where ***H*** is a reference vector field. The metric tensors of the reference and current states induced by *ψ*_*0*_ and *ψ* can be computed by differentiating these maps with respect to *ξ*^*1*^, *ξ*^*2*^, and *ξ*^*3*^ to obtain the convected basis vectors, and then computing the scalar products of these vectors. The resulting metric tensors will be in general a function of *ξ*^*3*^. To obtain a reduced theory, we sample these tensors at the mid-surface (*ξ*^*3*^ = 0). Furthermore, for simplicity and without loss of generality, we consider a reference configuration with constant thickness and lateral surfaces perpendicular to Γ_0_, in which case the reference vector field can be chosen as ***H*** = *h*_0_ ***N***. With these assumptions, a simple calculation shows that

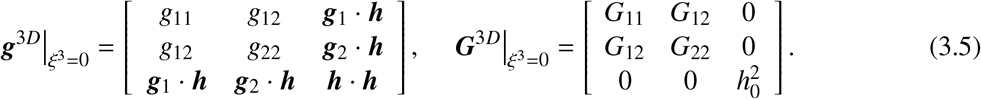

#### Thickness and cell incompressibility

Assuming that cells keep their volume constant [27], we impose local volumetric incompressibility in the epithelial shell with the condition 1 = *J*^3*D*^ = *g*^3*D*^/*G*^3*D*^, where *g*^3*D*^ and *G*^3*D*^ are the square roots of the determinants of the metric tensors in Eq. (3.5). Expanding the determinants by blocks, we have

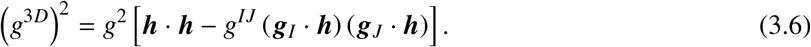

We note that *g*^*IJ*^ ( ***g***_*I*_· ***h***) ***g***_*J*_ is the tangential projection of ***h***, which satisfies that ***h*** = Π***h*** + (***h***· ***n***) ***n***. We thus have

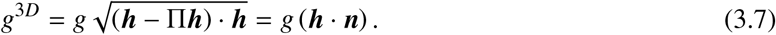

Since *G*^3*D*^ = *Gh*_0_, the local statement of 3D incompressibility of the tissue can be expressed as

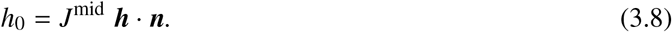

where we have introduced the notation *J*^mid^ = *g*/*G* to emphasize that this Jacobian determinant refers to the mid-surface of the shell. By introducing the scalar thickness of the tissue as *h*(ξ, *t*) = ***h***(ξ, *t*) · ***n***(ξ, *t*), this equation can be understood as imposing that the local volume, computed as the local tangential area times the thickness, remains constant.

#### Deformation of offset apical and basal surfaces

Given the map *φ* and the director field ***h***, we can define the ensemble of apical and basal faces of cells in the monolayer as offset surfaces 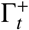 and 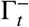 parametrized by

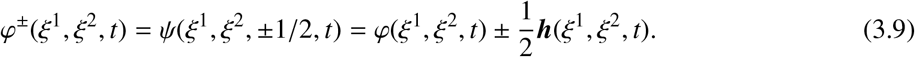

By doing this, we neglect the fact that apical and basal surfaces may bulge out, leading to apical and basal surfaces exhibiting cell-scale roughness. The above expression allows us to compute the corresponding metric tensors as

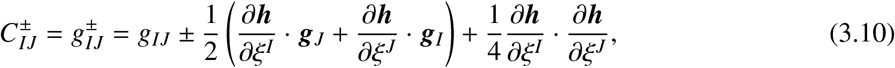

and

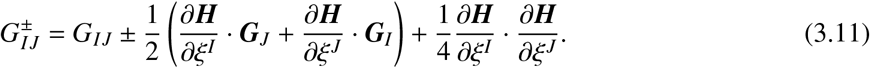

With these expressions, we can compute the right Cauchy-Green deformation tensor of the offset surfaces with mixed components as

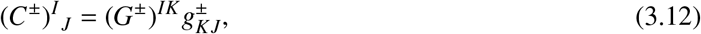

and introduce the Jacobian determinants of the offset surfaces as 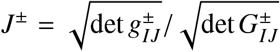 We can further compute the material time derivative of the metric tensors of offset surfaces as

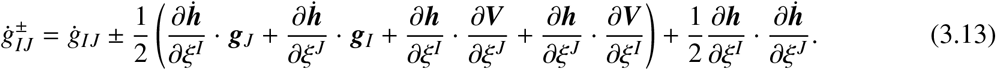

Since ***G***^±^ does not depend on time, this expression allows us to compute the rate-of-deformation as 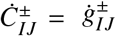, or with mixed components as 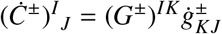.

#### Deformation of lateral surfaces

Consider now a lateral surface at position ***X* ∈** Γ_0_ and oriented in the reference configuration along a unit tangent vector ***M*** = *M*^*I*^***G***_*I*_ with *G*_*IJ*_ *M*^*I*^ *M*^*J*^ = 1 forming an angle *θ* ∈ (0, π) with ***G***_1_. We denote the normal vector to Γ_0_ by ***N***. Because we have chosen the reference configuration such that lateral faces are perpendicular to the mid-surface, it is natural to express the right Cauchy-Green deformation tensor of the lateral face in the orthonormal frame {***M, N***} . Recalling Eq. (3.5) and noting that the unit normal is expressed in convected basis induced by *ψ*_0_ as ***N*** = (1/*h*_0_)***G***_3_, we have

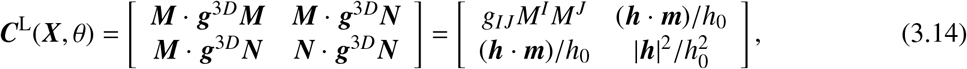

where we have introduced ***m*** = *M*^*I*^ ***g***_*I*_, a tangent vector to Γ_*t*_ not necessarily of unit length, see Fig. 3. We note that, because we use an orthonormal frame for the lateral surface,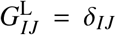. Therefore, the local change of lateral area can be expressed as 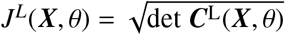. The strain rate then follows as

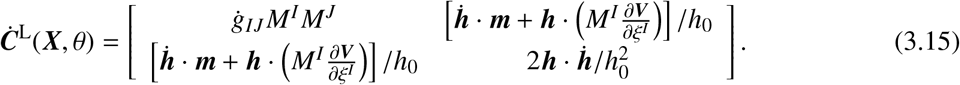

#### Coarse-grained description of the ensemble of lateral surfaces

In the continuum model, we do not track individual lateral cell-cell junctions but rather account for the effective behavior of an ensemble of lateral junctions within a *representative area element* (RAE) of tissue. Lateral junctions in the RAE oriented in different directions will deform differently given the overall deformation of the RAE. To account for the possibility that lateral junctions are not oriented isotropically, we coarse-grain the local ensemble of lateral junctions of a RAE located at point ***X* ∈** Γ_0_ with the angular distribution of the specific area of lateral junctions per tissue area in the reference state, which we denote by *f*_0_(***X***, θ). The total area of lateral junctions in the reference configuration of the tissue can be computed as 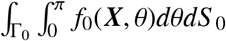The function *f*_0_(***X***, θ) thus embodies the anisotropy of the tissue and the aspect ratio of cells. For instance, for a “crystalline” tissue made of regular hexagonal prismatic cells with side length ℓ_0_ and height *h*_0_ and sides oriented at angles 0, *π*/3 and 2*π*/3, we can express the specific area distribution, independent of position, as

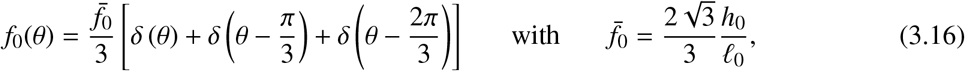

where *δ*(·) is the Dirac distribution. We can also define the angular distribution of the specific deformed area of lateral faces *f* (***X***, *θ*), which considering the local area changes in lateral faces and the tissue, can be related to *f*_0_(***X***, *θ*) by *f* (***X***, *θ*) =[ *J*^L^(***X***, *θ*)/*J*^mid^(***X***)]*f*_0_(***X***, *θ*).

### 3.2 Active viscoelastic model for the Cosserat epithelial shell

Integrating the ingredients of previous sections, we formulate next an effective Cosserat continuum model for the cell monolayer. Given the deformation map *φ* of the mid-surface and the director field ***h***, we can compute the deformation of the offset surfaces with Eqs. (3.10,3.11,3.12), and the deformation of a lateral face oriented along angle *θ* in Γ_0_ with Eq. (3.14). Lateral surfaces are distributed according to the angular distribution *f*_0_(*θ*). We introduce apical, basal and lateral density fields,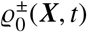 and 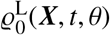 as well as the material metric tensor fields capturing viscoelastic relaxation G^±^(***X***, *t*) and G^L^(***X***, *t, θ*). Balance of mass for apical, basal and lateral faces is given by

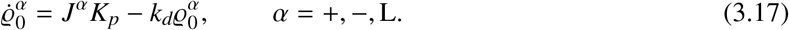

To write a coarse-grained tissue Rayleighian accounting for the Rayleighians of the ensemble of cell surfaces, we express the area element of apical and basal surfaces in the reference state in terms of that of the reference mid-surface, *dS* _0_ = *Gd*ξ^1^*d*ξ^2^. Defining 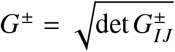 and since 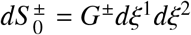, we have that 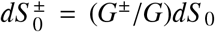. In particular, for a planar reference tissue,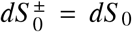. Similarly, by the very definition of *f*_0_(*θ*), we can express the area element of lateral surfaces oriented along *θ* as *f*_0_(*θ*)*dS* _0_. Thus, we can form the coarse-grained Rayleighian of the tissue collecting the contributions of all basal, apical and lateral surfaces as

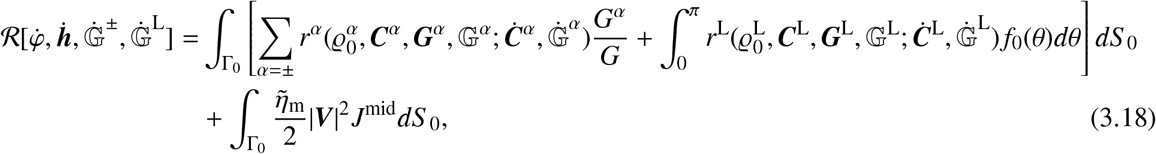

where the form of the Rayleighian density is given in Eq. (2.13). The first term in the first line accounts for the ensemble of apical and basal surfaces of the cell monolayer, the second term in the first line accounts for the ensemble of lateral surfaces, and the last integral models the rate-dependent friction of the medium surrounding the tissue. We note that this form of the medium dissipation is simplified to avoid the solution of the bulk hydrodynamic equations. The ± and L labels of the Rayleighian densities reflect the fact that the cortical mechanical properties modeled by functions Ψ, Φ and activity parameter *ξ* may be different in apical, basal and lateral faces.

To account for cellular incompressibility, we introduce the Lagrange multiplier field σ to enforce Eq. (3.8), and the constrained Rayleighian

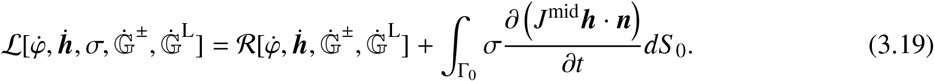

The governing equations of the theory are given by mass balance in Eqs. (3.17) and by the stationarity conditions of ℒ with respect to its arguments. The kinetic equations for the material metric tensors follow from the conditions 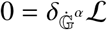 and can be expressed analogously to Eq. (2.14) as

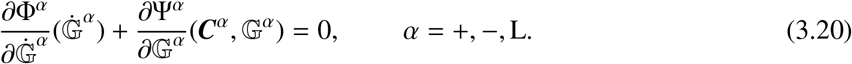

We note that for a continuous angular distribition *f*_0_(θ), Eqs. (3.17,3.20) for α = L represent a continuous family of evolution laws for density and material metric along each θ. The weak form of balance of linear momentum follows from directly from the stationarity condition 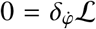. Analogously, the weak form of generalized force balance power-conjugate to 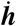 follows from the the stationarity condition 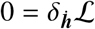. These weak equations are used directly in the finite element approximation of the theory.

## 4. Continuum Kirchhoff theory for epithelial shells

### 4.1. Kinematics under the Kirchhoff hypothesis

We develop next a theory akin to shell theories relying on the Kirchhoff hypothesis, according to which lines perpendicular to the mid-surface before deformation remain perpendicular after deformation [62]. This theory thus neglects through-the-thickness shears, an hypothesis usually valid for slender shells, which is not necessarily the case in epithelial sheets. This theory can be obtained from that in the previous section by requiring that the director field is normal to the surface, i.e. ***h*** = *h****n*** where 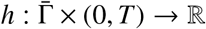 is a thickness scalar field. Consistent with the Kirchhoff hypothesis, we thus assume that lateral cellular faces are initially perpendicular to the tissue and remain perpendicular to Γ_*t*_ during deformation. See Fig. 4 for a graphical illustration of the Kirchhoff model.

**Figure 4:**
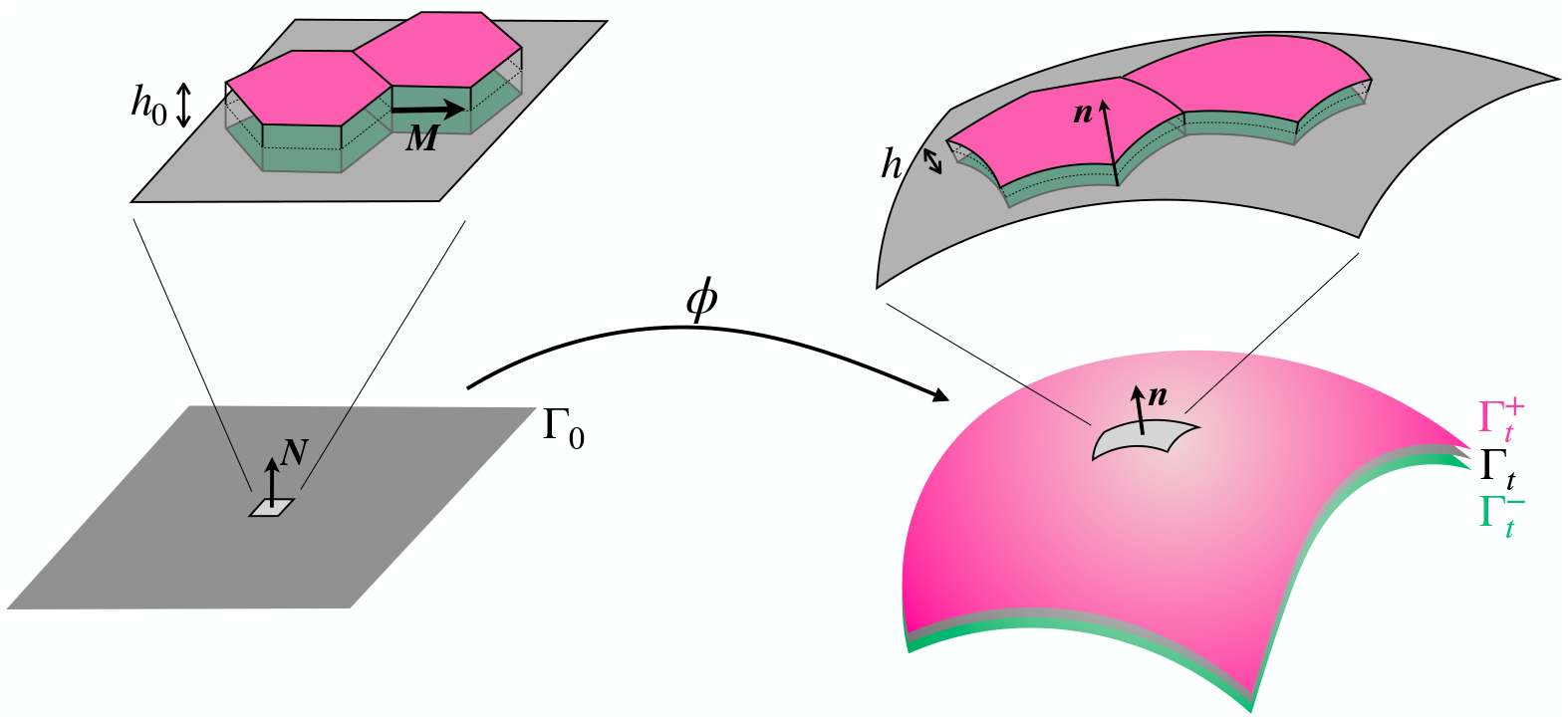
Illustration of the active bilayer model with the Kirchhoff hypothesis. For simplicity of illustration, we consider a planar reference configuration (left). We view the tissue as a continuum surface with mid-plane Γ_0_, microscopically composed of prismatic cells of height *h*_0_, with apical (pink) and basal (green) surfaces parallel to Γ_0_ and with lateral surfaces normal to Γ_0_ and oriented in the plane along directions given by the vector ***M***. In the deformed configuration (right), the mid-surface is Γ_*t*_. The offset surfaces 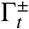 represent the ensemble of apical and basal surfaces. We assume that the lateral surfaces remain normal to the mid-surface Γ_*t*_.

#### Thickness and cell incompressibility

The local change of volume in the continuum model of the tissue can be approximated as the product of the local change of area of the mid-surface, *J*^mid^, and the local change in thickness. Cellular incompressibility then implies that *J*^mid^*h*/*h*_0_ = 1. Hence, we can express explicitly the thickness of the layer in terms of the Jacobian of the mid-surface as

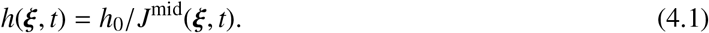

#### Deformation of offset apical and basal surfaces

Now, the parametrizations of the offset apical and basal surfaces, 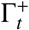 and 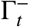, take the form

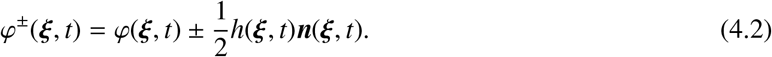

A direct calculation shows that

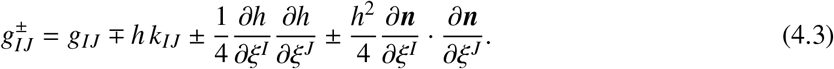

Recalling Eq. (4.1), we have *∂h*/*∂*ξ^*I*^ = − *h*_0_/(2*J*)*g*^*KL*^*∂g*_*KL*_/*∂*ξ^*I*^, and therefore it is clear that the metric tensor of the offset surfaces can be computed from the map of the mid-surface *φ* and its derivatives. In practice, and since the Kirchhoff theory may be pertinent to thin cell monolayers, it may be convenient to retain terms up to first-order in thickness

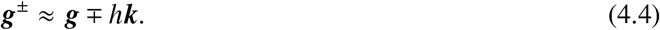

The same rationale can be applied to the reference state of the tissue to compute

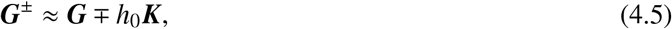

which for a planar reference state simplifies to ***G***^±^ = ***G***. As in the Cosserat theory, these expressions allow us to compute the right Cauchy-Green deformation tensor of the offset surfaces with mixed components as 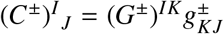 and the Jacobian determinants of the offset surfaces as 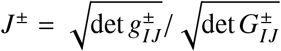. The material time derivative of the metric tensor, to first order in *h*, can be expressed as

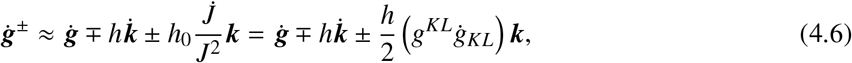

which allows us to compute the rate-of-deformation as 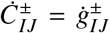, or with mixed components as 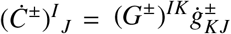.

#### Deformation of lateral surfaces

Consider a lateral surface aligned along the unit tangential vector in the reference configuration ***M*** forming an angle *θ* ∈ (0, *π*) with tangent vector ***G***_1_. Because we assume that lateral faces remain perpendicular to Γ_*t*_, the right Cauchy-Green deformation tensor is diagonal in the orthonormal frame {***M, N***} with components

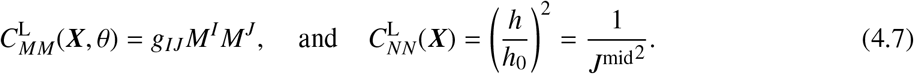

Thus, the deformation of a lateral junctions oriented along *θ* is characterized by

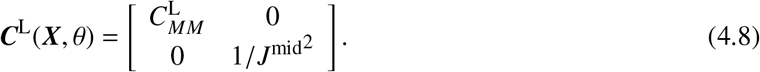

The local change of area of a lateral cell-cell junction along θ can therefore be expressed in terms of the deformation of the mid-surface as

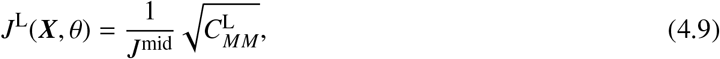

and the rate of deformation as

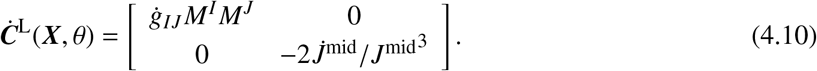

### 4.2 Active viscoelastic model for the Kirchhoff epithelial shell

Analogously to the Cosserat case, we are in a position to formulate an effective continuum model for the cell monolayer under the Kirchhoff hypothesis. Given the deformation map *φ* of the mid-surface, we compute the thickness of the bilayer with Eq. (4.1), the deformation of the offset surfaces with Eqs. (4.4,4.5,3.12), and the deformation of a lateral face oriented along angle θ in Γ_0_ according to Eq. (4.8). Following the same rationale as before, we can form an effective Rayleigian with the same form as that in Eq. (3.18), except that now it does not depend on the director field,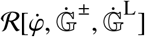. The arguments to obtain the evolution equations for the viscoelastic metric tensors and the weak form of the force balance equations are analogous, and the equations of balance of mass take the same form as in Eq. (3.17).

For the Kirchhoff theory, we further elaborate the Rayleighian by introducing Eqs. (4.6,4.10) for 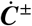 and 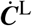 into Eq. (3.18). Keeping up to linear terms in *h*, a lengthy but otherwise direct manipulation leads to the following expression

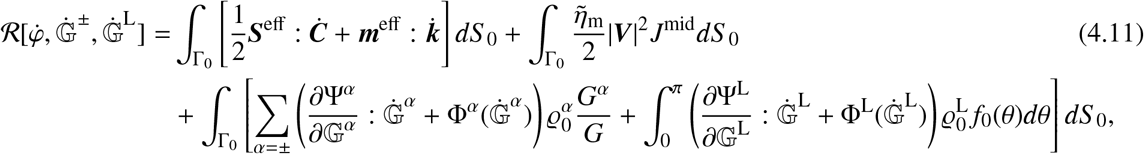

where the first line collects terms involving 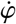, the second line terms involving rates of change of the material metric tensor fields, and we have introduced

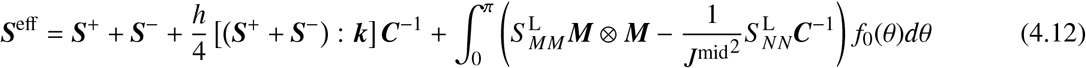

and

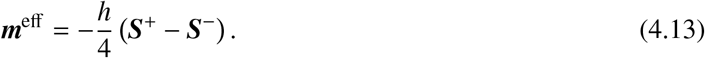

In these expressions, ***S***^±^ and ***S***^L^ denote the second Piola-Kirchhoff stress tensors of the apical/basal offset surfaces and of lateral surfaces, see Eq. (2.19),

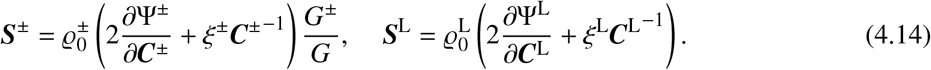

This re-elaboration of the Rayleighian allows us to identify ***S***^eff^ as the effective Lagrangian membrane stress tensor and ***m***^eff^ as the Lagrangian bending moment tensor for the homogenized model. Therefore, even though we have modeled the underlying cortex as a membrane-like surface that only supports in-plane stresses, the effective model is a shell-like surface that also supports bending moments, albeit emerging from localized surface tensions separated by a distance *h*. While the in-plane stress ***S***^eff^ depends on the sum of apical and basal membrane stresses and the net stress coming from lateral junctions (integral term in Eq. (4.12)), the bending moment tensor ***m***^eff^ depends only on the difference of apical and basal membrane stresses. These relations clearly show how in our model the effective stresses in the tissue do not result from phenomenological macroscopic constitutive relations but rather from the microscopic behavior of cellular surfaces modeled with an active viscoelastic gel.

In our model, the stress in the reference configuration (*C*^*I*^ _*J*_ = δ^*I*^ _*J*_) is not zero in general, even at steady-state when the elastic stresses have relaxed. To see this, consider for simplicity a planar reference tissue with an isotropic distribution of lateral junctions, that is 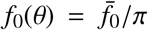 but the same result holds when it follows Eq. (3.16). In this situation, only the active stress contribution remains and is also isotropic. Particularizing Eq. (4.12), a direct calculation shows that the mean effective tension (trace ***S***^eff^)/2 in the reference configuration and at steady state takes the form

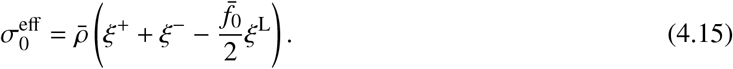

Because it is defined at the reference state, this expression is also valid for the Cosserat theory. It shows how the prestress of the tissue in the reference configuration depends on the aspect ratio parameter 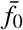 and on the apical, basal and lateral tensions.

## 5. Constitutive choices

To fix the ideas, we make specific choices for the material functions and parameters. For the angular distribution of lateral faces *f*_0_(θ), we discretize it as in Eq. (3.16), which can be interpreted either as describing of a crystalline tissue or as an approximation of a uniform distribution. Because this distribution is concentrated around three Dirac deltas, the resulting theory introduces only three families of lateral surfaces, and hence three fields of lateral material metric tensors, which we denote by G^α^(***X***, *t*) with α = 1, 2, 3. Likewise, we denote the Cauchy-Green deformation tensor of these three families of lateral faces by ***C***^α^ with α = 1, 2, 3. To simplify the implementation, we assume that the density *ρ* is constant. Physically, this is a reasonable approximation in a fast turnover regime relative to viscoelastic relaxation, in which case the density *ρ* reaches the constant steady-state value 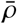 much faster than any other process. Even if this is not the case, calculations accounting for turnover show that with reasonable active gel parameters, this approximation may not affect much the out-of-equilibrium response of epithelial monolayers [34]. Consequently, the density per reference area in apical, basal and lateral surfaces can be computed as

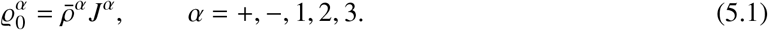

For the stored elastic energy, we consider a standard neohookean model of the form

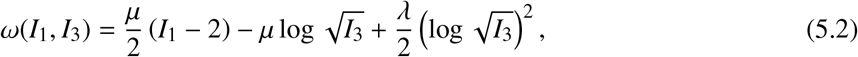

where *µ* and *λ* are 2D Lamé parameters of the cortical surface, and define the elastic potential per unit area as

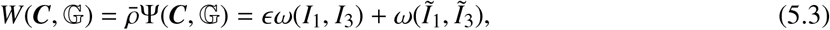

where ϵ controls the amount of residual elasticity of the cell surfaces that is not susceptible to viscoelastic relaxation. For the dissipation potential, we chose a quadratic potential of the form in Eq. (2.8) with **η**_1_ = 0 and we introduce the viscosity parameter 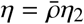. We also introduce the active tension parameter 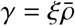.

In principle, the parameters 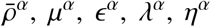 and *γ*^*α*^ can be different for apical, basal and lateral surfaces, *α* = +, −, 1, 2, 3. Again, to fix the ideas, we assume that ϵ is constant across surfaces, and that the other parameters of apical and basal surfaces are uniformly scaled from the reference parameters of lateral surfaces (*µ* = *µ*^L^, *λ* = *λ*^L^, **η** = **η**^*L*^, and *γ* = *γ*^L^) according to the dimensionless factors [26]

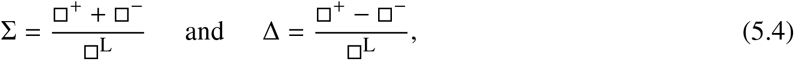

where □ stands for *µ, λ, *η**, or *γ*. With these choices and for an initially planar tissue, the Rayleighian of the tissue in the Kirchhoff theory can then be expressed as

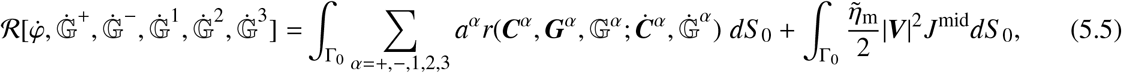

where 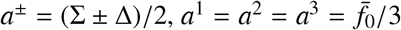, and

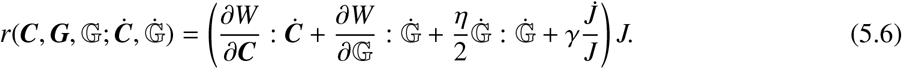

For the Cosserat theory, the constrained Rayleigian is

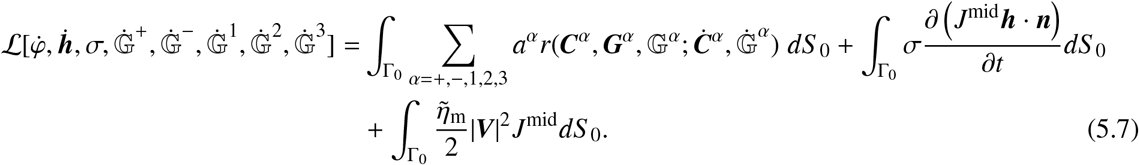

In summary, the mechanical parameters of the tissue in the bilayer model are the reference cortical properties (elastic coefficients *µ, λ* and *ϵ*, viscosity **η**, and active tension *γ*), the ratio of apicobasal to lateral mechanical properties Σ, the apicobasal asymmetry of mechanical properties Δ, and the parameter 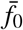 introduced in Eq. (3.16) encoding the aspect ratio of cells in the reference configuration. In addition, 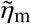 controls the friction of the tissue with the environment. For a system of typical linear dimension *L*, the following dimensionless quantities characterize the system: the ratio of Lamé parameters *π*_1_ = *λ*/*µ*, the normalized active tension (or elastocapillarity parameter) *π*_2_ = *γ*/*µ*, the normalized viscosity 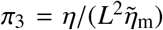, the slen-derness parameter *π*_4_ = *L*/*h*_0_, and the dimensionless parameters *ϵ*, Σ, Δ and 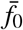. If the problem studied has a time-scale *τ*, then, an additional dimensionless number *π*_5_ = *τ*/ *τ*_ve_ arises from the comparison with the time of viscoelastic relaxation *τ*_ve_ = **η**/*µ*. We finally note that Eq. (4.15) shows that at an initially planar and isotropic reference configuration, the dimensionless steady-state prestress of the tissue is

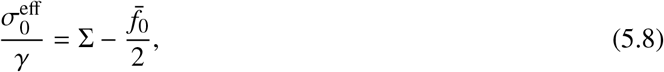

which suggests introducing the dimensionless prestress parameter

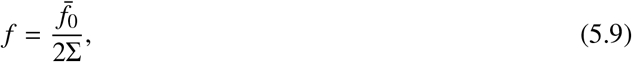

allowing us to distinguish between initially tense ( *f* < 1) and compressed ( *f* > 1) tissues.

## 6. Numerical implementation

The governing equations of the Cosserat and Kirchhoff theories of epithelial shells are highly nonlinear, and therefore we resort to numerical approximation. We use Galerkin finite elements for space discretization, and develop a time-discrete variational principle for time-marching.

### 6.1 Time discretization and incremental Onsager principle

For time-integration, we derive an implicit backward-Euler variational integrator by formulating an incremental Rayleighian functional, which is minimized with respect to the unknown state variables in the next time-step. This kind of structure-preserving integrator is interesting because it inherits qualitative properties of the time-continuous system [63, 41, 64], here the entropy production inequality in the absence of power input.

We consider a sequence of time instants *t*^1^, *t*^2^, … , *t*^n^, *t*^n+1^, …, and define the time-step size as Δ*t*^n+1^ = *t*^n+1^ − *t*^n^. The super-index on any time-dependent function denotes its time instant of evaluation. To define an incremental Rayleighian, we rewrite the Rayleighian density for a cortical surface in Eq. (2.13) as

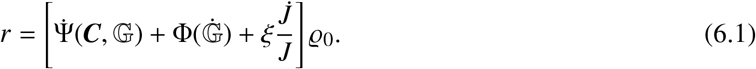

Approximating time derivatives with a first-order backward-Euler formula, e.g.

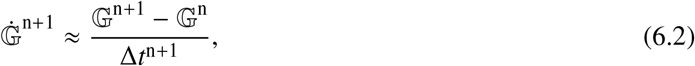

evaluating at *t* = *t*^n+1^ and multiplying by Δ*t*^n+1^, we define the incremental Rayleighian density as

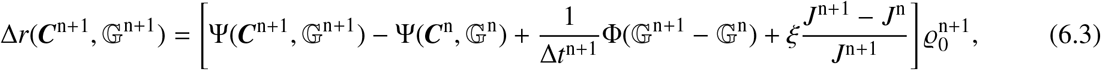

where we have emphasized the dependence of Δ*r* on the unknown variables in the next step. To obtain this expression, we have used the fact that 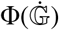is homogeneous of degree 2.

Adopting the limit of fast turnover as discussed in the previous chapter, we have 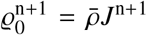, where 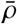 is the steady-state cortical density per actual area, assumed to be constant. Then, with the constitutive choices and the definitions introduced in Section 5., we can formulate the incremental Rayleighian density of cortical surfaces as

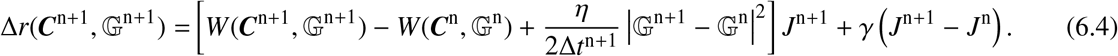

This Rayleighian density allows us to formulate the incremental Lagrangian of the Cosserat theory as

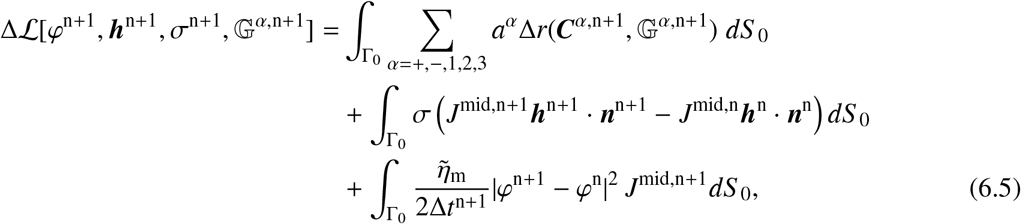

where we have used the same backward-Euler approximation for the time-derivative appearing in the volume constraint and for the Lagrangian velocity field. Analogously, we can formulate the incremental Rayleighian of the Kirchhoff theory as

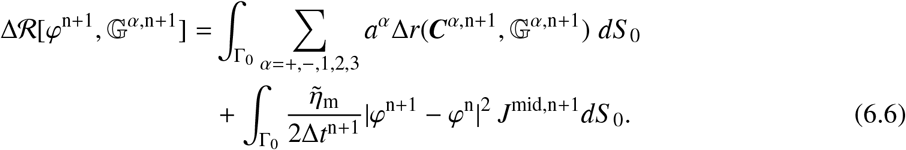

### 6.2 Space finite element discretization

The different nature of the Cosserat and Kirchhoff theories leads to different numerical discretizations. On the one hand, in the Cosserat theory, the description of the kinematic state of the system involves six degrees of freedom (*φ* and ***h***), and an additional Lagrange multiplier field (*σ*) is needed to enforce the incompressibility constraint, Eq. (3.8). Instead, in the Kirchhoff theory, deformation is described by only three degrees of freedom (*φ*). On the other hand, while the formulation of the Rayleighian in the Cosserat theory only involves up to first order derivatives of *φ* and ***h***, and therefore straightforward *C*^0^ finite elements can be used, the Kirchhoff Rayleighian involves second derivatives to compute the curvature, see Eqs. (2.2,4.6). Consequently, the Kirchhoff theory demands a more sophisticated numerical treatment. Here, we choose to approximate *φ* in the Kirchhoff theory with subdivision finite elements, a spline technique supported on an unstructured triangulation of the surface and leading to *C*^1^ surface approximations [65, 41]. In the Cosserat theory, we use subdivision finite elements and *C*^0^ linear finite elements on quadrilateral meshes, depending on the example. We also note that the Rayleighian functional does not involve space derivatives of the metric tensors G^*α*^(*t*) with *α* = +,− , 1, 2, 3. Consequently, these 5 fields do not need to be interpolated and can be directly defined and evolved locally, at the level of each quadrature point of each element.

To discretize the mid-surface of the shell with finite elements, we choose as parametric domain 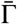 introduced in Section 2.1 the parent finite element, e.g. a triangle described with barycentric coordinates, and approximate the fields *φ*_0_(*ξ*), and *φ*(*ξ, t*) with functions defined piecewise over the elements as

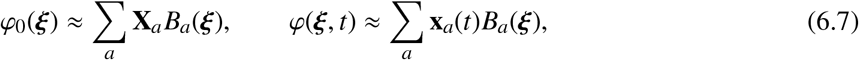

where *B*_*a*_(*ξ*) are the basis functions supported on a surface mesh, and the index *a* runs over the nodes contributing to the approximation over the reference element. For the Cosserat theory, we approximate the fields ***H***(*ξ*), ***h***(*ξ, t*), and *σ*(*ξ, t*) analogously as

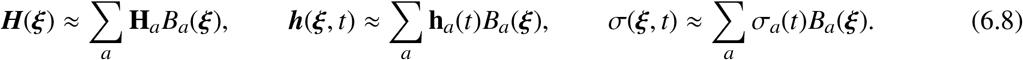

### 6.3 Fully discrete governing equations

The fully discrete equations to move forward in time from *t*^n^ to *t*^n+1^ are obtained in the Cosserat theory by plugging the approximations in Eqs. (6.7,6.8) into the time-incremental constrained Rayleighian in Eq. (6.5), and then extremizing the functional with respect to the nodal degrees of freedom 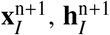 and 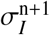 and with respect to the material metric tensors at each quadrature point. To simplify the implementation and reduce the size of the resulting nonlinear system of equations, we choose to solve the nodal degrees of freedom and the material metric tensors in a staggered way. Thus, denoting the total number of nodes in the mesh by *N* and the total number of quadrature points by *Q*, in each time-step we need to solve 1 nonlinear system of 7*N* unknowns (position, director and Lagrange multiplier) and 5*Q* nonlinear systems with 3 unknowns (metric tensors). We solve these systems using Newton’s method. We adapt the time-step with two criteria. First, we set a maximum step-size that we are willing to accept to follow the dynamics with sufficient accuracy. Second, we increase/decrease the time-step using a multiplicative factor if the number of Newton iterations is lower/higher than a prescribed threshold.

## 7. Results

### 7.1 Buckling by deflation of an epithelial blister: comparison with a 3D cellular model

To verify our continuum theory, we perform a direct comparison of simulations based on the coarse-grained continuum model with simulations of the tissue explicitly modeling each cell as a collection of cortical surfaces enclosing constant volumes [34], where each cellular surface is modeled as described in Section 2.. Because the tissue is quite slender, we consider first the Kirchhoff continuum theory. We report all the parameters used in the simulations in this work in Table 1.

**Table 1.** Model parameters

| | $\lambda/\mu$ | $\gamma/\mu$ | $\eta/(L^2\tilde{\eta}_m)$ | $L/h_0$ | $\Sigma$ | $\Delta$ | $f$ | $\epsilon$ |
| --- | --- | --- | --- | --- | --- | --- | --- | --- |
| Fig. 5 | 1 | 0.5 | 218 | 26 | 2 | 0 | 0.04; 0.19; 1.5 | 0 |
| Fig. 6 | 1 | 0.03 | 1000 | 50 | 2 | 0, 0.2, 0.4 | 0.7 | 0 |
| Fig. 7 | 1 | 0.015 | 1000 | 25, 50, 80 | 2 | 0.04 to 0.12 | 1 | 0.01 |
| Fig. 8 | 1 | 0.1 | 1000 | 10 | 5 | 0 to 3 | 1.5 | 0 |

As a representative example of the complex and time-dependent epithelial mechanics, we study a situation leading to buckling dynamics. We consider an initially planar cell monolayer attached to a substrate except in a circular region. We consider as the typical length-scale of the problem *L* the diameter of the unattached footprint. Upon pressurization, the free-standing part of the tissue inflates into a nearly spherical cap geometry [19]. See Fig. 5(a) for an illustration. We condition the tissue by a long inflation time (≫ *τ*_ve_) so that the cortex of each cellular surface reaches its steady state in the inflated configuration. If the time of deflation *τ*_d_ is small enough compared to the viscoelastic relaxation time *τ*_ve_, the tissue experiences compression and buckles, Fig. 5(b). This problem is extensively studied both computationally and experimentally elsewhere [66].

**Figure 5:**
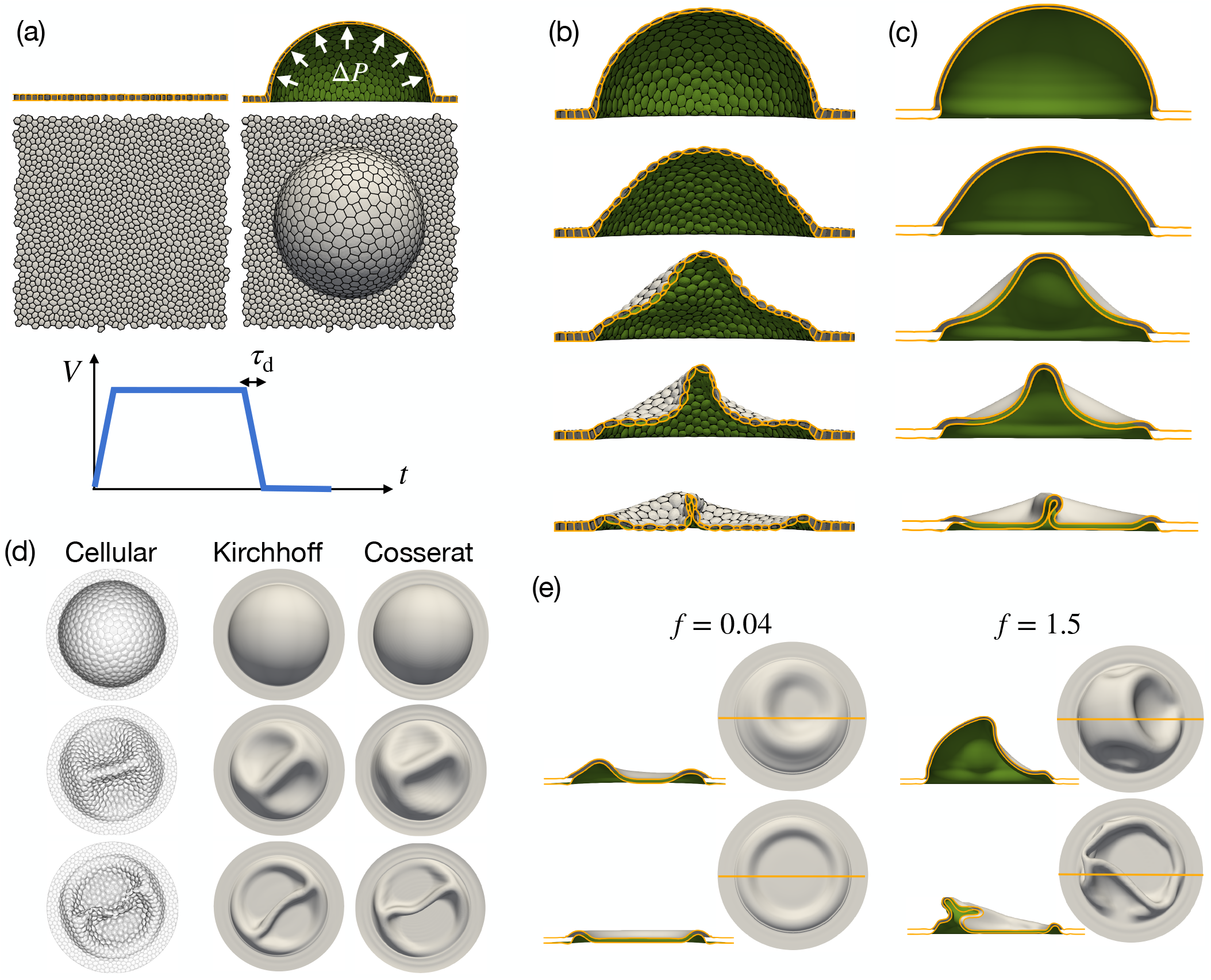
Comparison of the proposed continuum shell theory and a 3D cellular model based on active gels to study the buckling dynamics induced by the rapid deflation of an epithelial shell. (a) Illustration of the setup. Mimicking adhesion patterning, an initially planar tissue is attached to the substrate outside of a detached circular footprint. The volume enclosed between the tissue and the substrate is then increased and held for a long-enough time to achieve full viscoelastic relaxation. Then, it is reduced within the timescale *τ*_d_ = 0.6 *τ*_ve_. (b) Cross-section of the epithelial blister at selected instants during deflation for the 3D cellular model, and (c) corresponding cross-sections for the proposed Kirchhoff epithelial shell theory. See also Movie 1. (d) Top view of the apical surfaces at selected instants during deflation for the cellular model and for the Kirchhoff and Cosserat epithelial shell models. (e) Effect of the prestress parameter *f* on the buckling dynamics. This parameter is *f* = 0.19 in (a-d).

The direct comparison of the buckled morphologies at selected instants obtained with the two independently run simulations, one explicitly modeling each individual cell and one based on the continuum Kirchhoff theory proposed here, is shown in Fig. 5(b,c). For post-processing purposes, in the continuum simulations we show the apical and basal offset surfaces although computationally we only solve for the mid-surface. The figure shows that the bilayer continuum model disregards geometric details at the cellular scale, such as the bulging of apical and basal surfaces, but captures the overall shape of the buckling pattern remarkably well. The computational complexity of the continuum model is much smaller than that of the 3D cellular model. For instance, the computational mesh of the model in Fig. 5(b) involves over 111,000 nodes, whereas that in Fig. 5(c) about 2,000 nodes. We finally compare the Kirchhoff and the Cosserat epithelial shell simulations for this problem, finding a very good agreement, Fig. 5(d), as expected for such a relatively thin epithelial monolayer.

These results show that for this problem it is not essential to retain cellular resolution to understand the buckling dynamics of epithelial sheets. Furthermore, it allows us to identify the key parameters that survive the coarse-graining procedure. Indeed, besides the parameters characterizing the actomyosin gel and the slenderness, our continuum model shows that the tissue architecture can be summarized by the prestress parameter 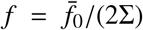, see Eq. (5.8). To illustrate the role of this parameter, we compare in Fig. 5(e) the baseline results ( *f* = 0.19), with those when *f* = 0.04, corresponding to a very tense tissue,which exhibits delayed buckling and small final excess area because prestress offsets to a large degree the viscoelastic compressive stress during deflation. Instead, when *f* = 1.5 > 1, the reference planar state is under compression and prestress at full inflation is low, and accordingly we observe early buckling and a larger excess area.

### 7.2 Effect of apicobasal asymmetry

#### Epithelial sheets suspended between supports

We examine next the ability of the proposed theories to model apicobasal asymmetry. We study the effect of apicobasal asymmetry in freestanding tissues attached to supports, simulating the experiments in [15]. We consider a rectangular prestressed tissue ( *f* = 0.7 < 1), constrain the displacements at two opposing ends and relax the system, modeled with the Cosserat theory. In the absence of apicobasal asymmetry (Δ = 0), the tissue contracts laterally, exhibiting curved free edges but remaining planar. Apicobasal asymmetry leads to steady-states exhibiting tissue curling largely localized at the free edge, Fig. 6(a), in close agreement with the experiments in [15]. Also in agreement with these experiments, we find that tissue stretching leads to partial uncurling of the free edge.

**Figure 6:**
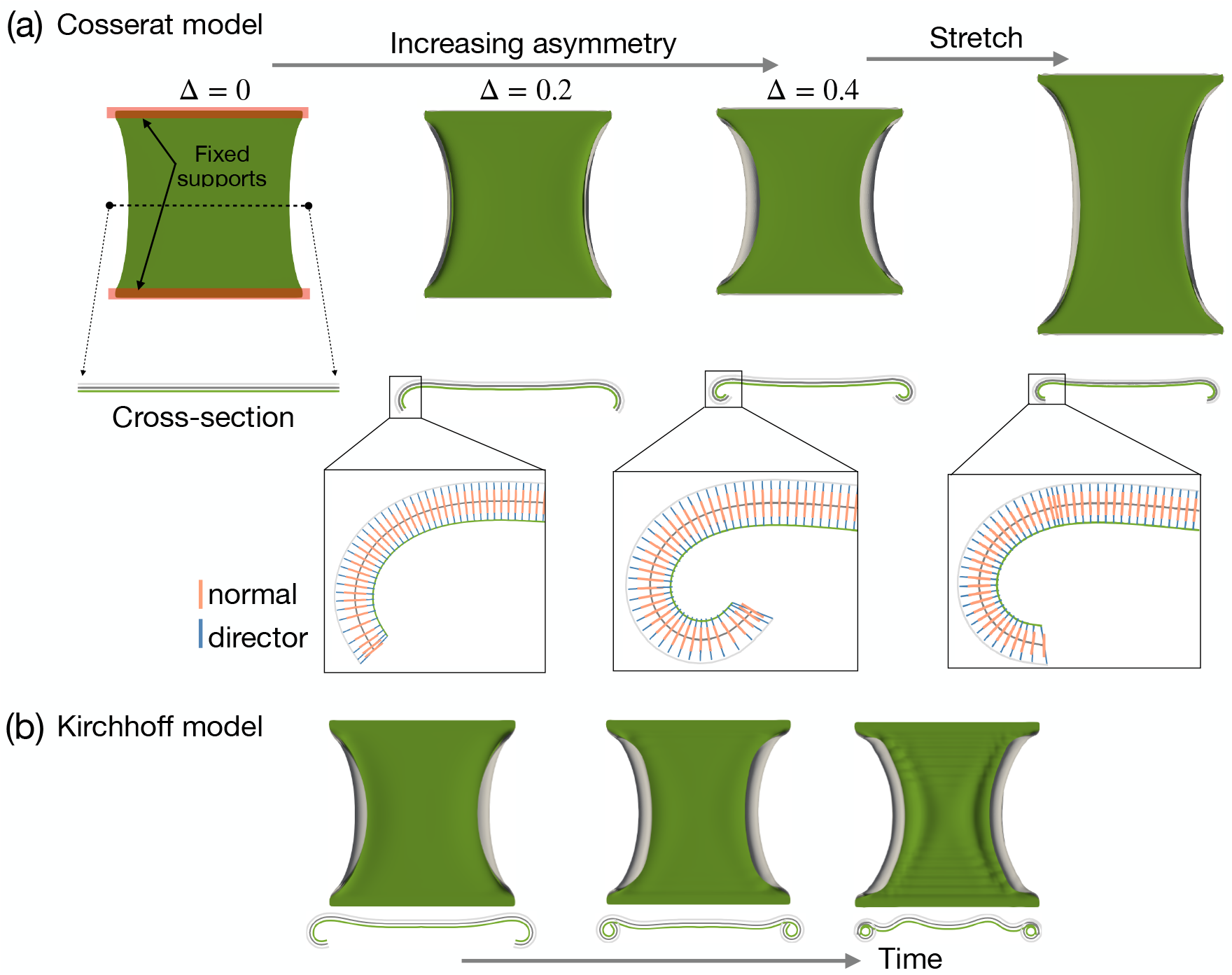
Prestressed tissue with two ends fixed. (a) Steady-states according to the Cosserat epithelial shell theory for increasing apicobasal asymmetry, and following a longitudinal stretch. See also Movie 2. The zoomed images at the bottom show the apical, middle, and basal surfaces, along with a sampling of the normal and director fields. The difference between the normal to the surface and the director field is most noticeable at the free-edges. (b) Snapshots of a simulation with the Kirchhoff epithelial shell theory for Δ = 0.4, which does not attain a steady-state.

This epithelial sheet being quite slender, we then examine the same problem using the Kirchhoff theory of epithelial shells. Unlike the problem of dome buckling, here the Kichhoff model fails to capture the mechanics of edge curling, and simulations fail to reach a steady-state with an increasingly intricate deformation until lack of convergence of the nonlinear solver, Fig. 6(b). The lack of a steady-state in the Kirchhoff theory can be understood from Eqs. (4.13,4.14). Indeed, at long times, the in-plane tensions of the apical and basal surfaces reduce to the asymmetrical active tensions, which generate bending moments that cannot be balanced by moments coming from any other source, and consequently, the system cannot reach equilibrium. In contrast the Cosserat model is able to generate balancing bending moments by tilting lateral surfaces; tilting introduces a tangential component of lateral tension acting with opposite sign along apical and basal surfaces, therefore leading to a bending moment that can balance that in Eq. (4.13). Consistent with this rationale, a close examination of the director field shows that it significantly deviates from the normal direction close to the free-edge, Fig. 6(a).

#### Floating disk

Even though cell monolayers are rarely completely free-standing, we examine the hypothetical situation of an unconstrained disc-shaped tissue. We consider the initial configuration to be tension-free ( *f* = 1) with finite apicobasal asymmetry Δ ≠ 0. As a background, we refer the reader to a related system consisting of two glued elastic sheets, with one of the sheets undergoing isotropic expansion or contraction [67]. Such an elastic bilayer typically exhibits spherical deformations at low asymmetry or slenderness, consistent with the isotropic nature of the underlying spontaneous curvature. However, at sufficiently large asymmetries, the stretching deformations required by Gauss’ theorem to change the Gaussian curvature from zero (initial planar state) to positive (spherical shape) trigger a transition to a nearly isometric curling of the disc into a cylindrical shape.

With the Kirchhoff theory, our simulations initially bend into spherical cap configurations, which depending on the parameters transits to a cylindrical configuration. However, in agreement with the discussion in the previous example, none of the simulations based on the Kirchhoff theory achieve a steady-state. When the transient shape before viscoelastic relaxation takes place is cylindrical, then the system evolves towards progressively tighter scrolls. Instead, when the transient shape is spherical, the system evolves towards intricate geometries, Fig. 7(a). As before, in the Cosserat model, tilt generates bending moments that can balance those resulting from apicobasal asymmetry and the system reaches a steady state. However, even for small asymmetries, the resulting equilibrium shape is a tight scroll, that exhibits self-intersection because we did not implement a contact algorithm.

**Figure 7:**
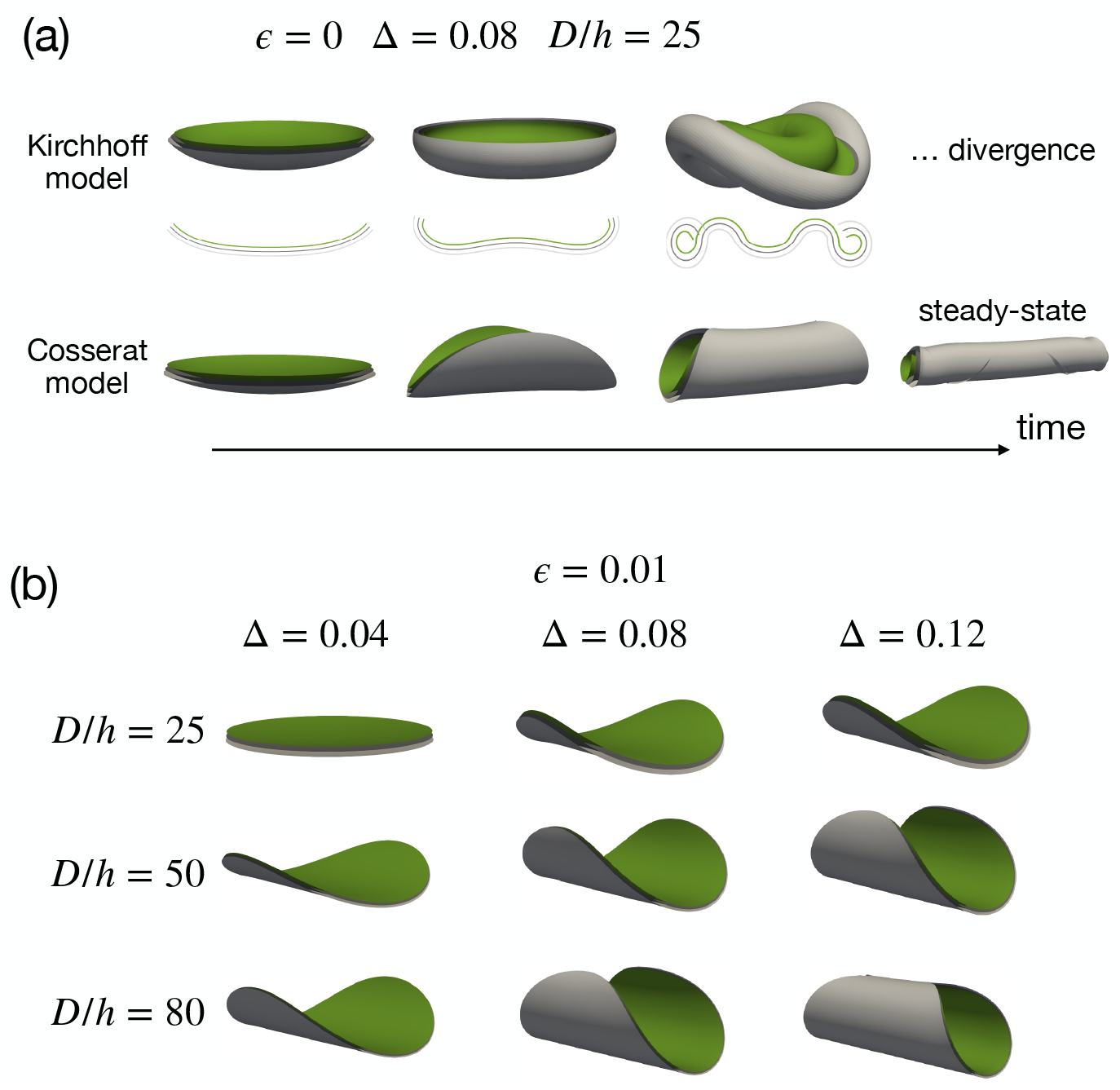
Effect of tension asymmetry (Δ ≠ 0) on free-standing discs. (a) Dynamics of a free-standing disk-like tissue without residual elasticity (*ϵ* = 0) simulated with the Kirchhoff model (top), which does not reach a steady state, and with the Cosserat model (bottom). (b) Steady-state configurations for different apicobasal asymmetry and slenderness parameters simulated with the Kirchhoff theory and a small value of the residual elasticity parameter.

An alternative way of introducing a physical mechanism to balance active moments arising from apicobasal asymmetry is to consider a finite amount of residual elasticity (*ϵ* > 0), in which case our simulations do reach a steady-state, Fig. 6(b). With a small amount of residual elasticity, the steady-state phenomenology of the Kirchhoff model is similar to that of elastic bilayers [67], with cylindrical steady-states being favored as both slenderness and asymmetry increase.

#### Wrinkling of compressed and asymmetrical epithelial sheets

We conclude by examining the effect of apicobasal asymmetry on tissue buckling. This problem has been studied by [26] with a 1D beam-like continuum model accounting for tilt and coarse-graining a 2D lateral vertex model. It hence serves as a verification of our Cosserat theory.

When a free-standing and symmetric epithelial monolayer is uniaxially compressed to a sufficient amount, we expect compressive tensions to develop at steady-state, and since the tissue is a slender structure, we expect Euler buckling [14, 34]. We consider a tissue with compressive prestress ( *f* = 1.5 > 1), and relax it under periodic boundary conditions. To favor uniaxial buckling, we consider a very thin tissue strip. As expected, the tissue exhibits Euler buckling with the longest wavelength fitting in the computational periodic domain, Fig. 8(a). The tissue being symmetric (Δ = 0), buckling takes place upwards or downwards with equal probability. When basal contractility is slightly increased relative to apical contractility (Δ = 0.25, 0.5), the asymmetry breaks the top-down symmetry of the buckled configuration and hints a buckling mode with higher wavenumber. Beyond a threshold of asymmetry, the system transitions to a wrinkled state with decreasing wavelength as asymmetry is increased (Δ = 0.75, 1.5), in agreement with the results reported by [26].

**Figure 8:**
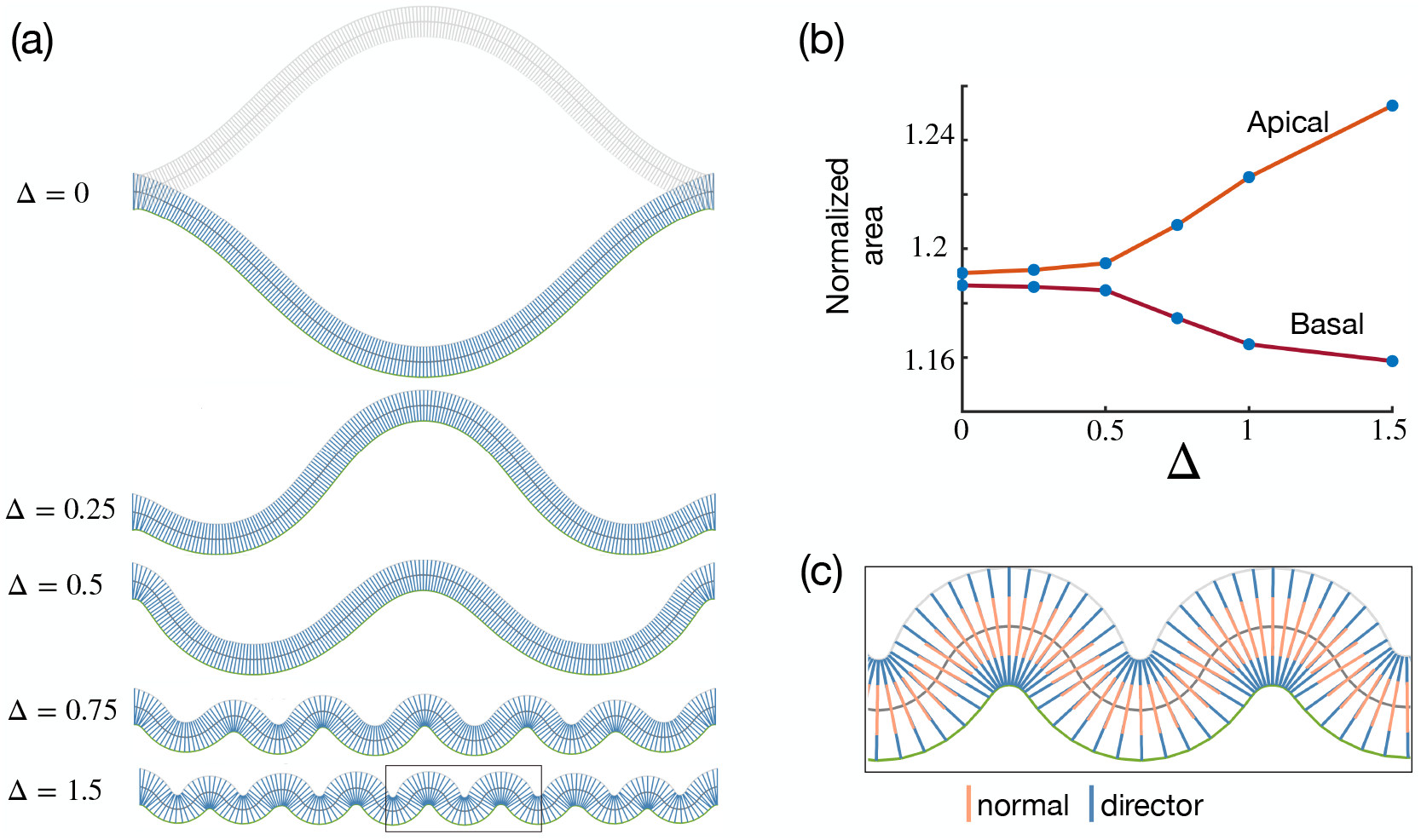
Wrinkling of free-standing tissues induced by apicobasal asymmetry. (a) Steady-state configuration for a tissue strip with periodic boundary conditions as the asymmetry parameter increases. See Movie 3 for the dynamics. (b) Total apical and basal areas normalized by the projected area of the tissue in the periodic box. (c) Detail of the apical, middle, and basal surfaces, along with a sampling of the normal and director fields, for a wrinkled configuration.

Conventionally, wrinkling arises when a thin compressed structure is confined by a surrounding medium, as classically studied by [68], with a wavelength set by a competition between the elastic deformation of the thin structure and that of the surrounding medium. Here, instead, the tissue wrinkles in the absence of a surrounding medium. The analysis by [26] shows that beyond a threshold Δ, a “phantom” substrate appears as a result of tension asymmetry and compression. To physically understand this unconventional wrinkling phenomenon, we examine the apical and basal areas as Δ increases, finding that the wrinkled configurations reduce the area of the basal surface at the expense of increasing the apical area, Fig. 8(b), which may be energetically favorable if the basal tension is significantly larger than the apical one. As in the previous examples, the Kirchhoff theory fails to reach a steady state, suggesting the need for tilt when studying wrinkling induced by apicobasal asymmetry. Accordingly, our simulations with the Cosserat theory show that the director field significantly deviates from the normal, Fig. 8(c). Thus, epithelial shells develop a wrinkling mode associated with apicobasal asymmetry and mobilizing the tilt of lateral surfaces. This example further highlights the limitations of conventional elastic shell theories to capture the mechanics of epithelial shells.

Taken together, our simulations show that the Kirchhoff theory provides a good approximation to nonlinear epithelial mechanics when the tissue is slender and symmetric. However, in the presence of apicobasal tension asymmetries, this theory fails to converge to steady-states because it lacks a physical mechanism to balance the active bending moments resulting from asymmetry. This deficiency is not present in the Cosserat theory, where tilt provides a mechanism to generate bending moments arising from tension in lateral faces.

## 8. Conclusions

We have presented a fully nonlinear, active, and viscoelastic theory of epithelial shells that, rather than relying on phenomenological constitutive relations, homogenizes a microscopic 3D-vertex-like model accounting for individual cell surfaces. In turn, these cell surfaces are described as active gel patches. The proposed approach results in a continuum surface model that cannot be directly mapped to conventional shell theories, and that captures the net effect of an ensemble of active viscoelastic active gel surfaces undergoing turnover. Our theory can account for the geometric and mechanical anisotropy of lateral junctions, as well as for the apicobasal asymmetry of cortical mechanical properties. The continuum effective model presents several advantages with respect to a detailed cellular model; it leads to much more efficient simulations, it is amenable to mathematical analysis, and it clearly identifies the essential model parameters. For instance, we have shown that in the present context, the tissue architecture is coarse-grained at the continuum scale in terms of the function *f*_0_(*θ*) measuring the specific area of lateral surfaces along different orientations.

The key idea of our approach is to relate the deformation of cellular surfaces (apical, basal and lateral) to that of the mid-surface. We describe the ensembles of apical and basal surfaces as offset surfaces of the mid-surface of the tissue, and adjust the thickness of the resulting bilayer according to the hypothesis of cellular incompressibility. We have presented two flavors of the theory, a Kirchhoff epithelial shell theory, where lateral junctions remain perpendicular to the mid-surface, and a Cosserat theory where lateral junctions can tilt. The proposed coarse-graining procedure is straightforward thanks to a variational formalism for the dynamics of an active gel. In this formalism, the governing equations for the microscopic model minimize a Rayleighian functional accounting for viscoelasticity and active contraction of each cellular surface. By introducing the kinematic relations between cellular and continuum deformations, we obtain an effective Rayleighian functional in terms of continuum fields, from which the continuum governing equations and finite element equations can be derived. We have verified our model by comparing 3D cellular model simulations with the finite element discretization of our theory, obtaining a remarkable agreement and finding good agreement between the Kirchhoff and the Cosserat theories. We have further examined the effect of apicobasal asymmetry on free-standing tissues, finding that the tilt emerging from the Cosserat theory is essential to obtain stable steady-states.

The present work can be expanded in many ways. Here, we have focused on the actomyosin cortex, but thanks to the generality of the coarse-graining approach, the underlying microscopic model may include additional mechanical elements such as apical belts, intermediate filaments, or models for adhesion dynamics. It can also be coupled to deformable substrates, or adapted to plant cellular tissues. Our theory focuses on situations where tissue deformation results from cellular deformations. It thus assumes that the architecture of the tissue does not evolve. However, tissue deformation can also be the result of cell divisions, cell extrusion, or junctional rearrangements [10]. Interestingly, because our work shows that tissue architecture is described at the continuum scale by the function *f*_0_(*θ*), it also provides a continuum framework to model the evolution of tissue architecture within the context of irreversible thermodynamics by viewing *f*_0_(*θ, t*) as a dynamical variable. Another interesting avenue for further research is developing continuum models that couple mechanics and signaling. Indeed, mechanical properties such as contractility or adhesion dynamics are controlled by morhogens, which in turn are also mechanosensitive. Previous work has studied effective continuum models for morphogen transport starting from cellular and sub-cellular mechanisms [69]. The combination of this kind of signaling model with continuum mechanical models such as those presented here could provide a powerful approach to understand epithelial tissue mechanobiology.

## Acknowledgements

This work was supported by the European Research Council (CoG-681434), the Spanish Ministry of Science, Innovation and Universities and the Spanish State Research Agency MICIU/AEI/10.13039/501100011033 (PID2022-142178NB-I00), and the Generalitat de Catalunya (2021-SGR-01049 and ICREA Academia prize for excellence in research to MA). IBEC is recipient of a Severo Ochoa Award of Excellence.

